# Telomere relocalization to the nuclear pore complex in response to replication stress

**DOI:** 10.1101/2020.01.31.929059

**Authors:** Alexandra M Pinzaru, Noa Lamm, Mike al-Kareh, Eros Lazzerini-Denchi, Anthony J Cesare, Agnel Sfeir

## Abstract

Mutations in the telomere binding protein, POT1 are associated with solid tumors and leukemias. POT1 alterations cause rapid telomere elongation, ATR kinase activation, telomere fragility, and accelerated tumor development. Here, we investigated the impact of mutant POT1 alleles through complementary genetic and proteomic approaches based on CRISPR-interference and biotin-based proximity labelling, respectively. These screens revealed that replication stress is a major vulnerability in cells expressing mutant POT1 and manifest in increased mitotic DNA synthesis (MiDAS) at telomeres. Our study also unveiled a role for the nuclear pore complex (NPC) in resolving replication defects at telomeres. Depletion of NPC subunits in the context of POT1 dysfunction increased DNA damage signaling and telomere fragility. Furthermore, we observed telomere repositioning to the nuclear periphery driven by nuclear F-actin polymerization in cells with POT1 mutations. In conclusion, our study establishes that relocalization of dysfunctional telomeres to the nuclear periphery is critical to preserve telomere repeat integrity.

## Introduction

Mammalian telomeres are composed of tracts of duplex TTAGGG repeats that are extended by the telomerase reverse transcriptase (Greider and Blackburn 1985). Telomere repeats are bound by a six-subunit complex – termed shelterin – that prevents chromosome ends from being recognized as DNA double-strand breaks (DSBs)(de Lange 2005). Shelterin is composed of TRF1 and TRF2 that bind to the duplex region of the telomeres and recruit RAP1, TIN2, TPP1, and the single strand DNA (ssDNA) binding protein, POT1.

Protection of telomeres 1 (POT1) contains two N-terminal oligonucleotide / oligosaccharide binding (OB) fold domains that bind to (T)TAGGGTTAG sequence present at the 3’ overhang or internally within exposed segments of the telomere (Baumann and Cech 2001; Lei et al. 2004). Despite its sub-nanomolar affinity to single-stranded telomere DNA, POT1 relies on its interaction with TPP1 to be recruited to telomeres (Liu et al. 2004; Hockemeyer et al. 2007; Nandakumar et al. 2010) and the POT1-TPP1 heterodimer is anchored to the rest of the shelterin complex through TIN2 (Liu et al. 2004; Ye et al. 2004; Takai et al. 2011). POT1 binding to the 3’ overhang occludes telomerase from accessing its substrate and negatively regulates telomere length (Loayza and De Lange 2003). POT1 also plays a key role in telomere protection. Knockdown of POT1 in human cells induces DNA damage signaling at telomeres (Veldman et al. 2004; Hockemeyer et al. 2005). Rodent telomeres contain two closely related POT1 proteins, POT1a and POT1b. Simultaneous deletion of both genes from mouse cells leads to RPA loading, activation of ATR signaling, and reduced cellular proliferation (Hockemeyer et al. 2006; Wu et al. 2006; Denchi and de Lange 2007; Gong and de Lange 2010).

Sequencing of cancer genomes has identified recurrent mutations in *POT1*. Sporadic missense and nonsense mutations were first reported in chronic lymphocytic leukemia (CLL) (Quesada et al. 2011; Ramsay et al. 2013) and later found in parathyroid adenoma (Newey et al. 2012), mantle cell lymphoma (Zhang et al. 2014), and adult T cell leukemia/lymphoma (Kataoka et al. 2015). Additionally, germline mutations in *POT1* were associated with familial cancers, including melanoma (Robles-Espinoza et al. 2014; Shi et al. 2014), and glioma (Bainbridge et al. 2015). *POT1* mutations are enriched within its OB fold domains and many alterations disrupt the binding of POT1 to telomere ssDNA in vitro (Calvete et al. 2015; Pinzaru et al. 2016). Consistent with cancer-associated *POT1* mutations causing telomere deprotection, expression of *POT1* OB-fold mutations in human cells results in rapid telomere elongation, telomere fragility, and ATR activation (Ramsay et al. 2013; Shi et al. 2014; Calvete et al. 2015; Pinzaru et al. 2016).

Telomere DNA is prone to forming stable secondary structures (Parkinson et al. 2002) that can impede replication fork progression and lead to telomere fragility (Martinez et al. 2009; Sfeir et al. 2009). Several telomere-associated factors assist in the synthesis of telomere DNA and prevent replication stress. For example, the shelterin subunit TRF1 recruits two helicases, RTEL1 and BLM, to unwind secondary structures and prevent stalling of the replisome as it copies telomeres (Sfeir et al. 2009; Vannier et al. 2012; Zimmermann et al. 2014). On the other hand, the polymerase-α primase accessory complex, CST (CTC1-STN1-TEN1), counteracts fork stalling by facilitating re-priming of DNA synthesis (Stewart et al. 2012; Wang et al. 2012; Kasbek et al. 2013). DNA combing analyses have shown that telomeres frequently undergo fork stalling events in cells expressing mutant POT1 (Pinzaru et al. 2016). The mechanistic basis by which *POT1* mutations compromise fork progression remains unknown, yet genetic evidence suggests that they act in the same pathway as the CST complex (Pinzaru et al. 2016).

Studies in *S. cerevisiae* revealed that eroded telomeres are targeted to the nuclear pore complex (NPC) (Khadaroo et al. 2009). Enrichment of telomeres at the NPC in yeast is driven by increased SUMOylation of RPA by Slx5-Slx8 (Churikov et al. 2016) and enhances the formation of type II survivors (Khadaroo et al. 2009). In mammalian cells, telomeres were shown to be transiently tethered to the nuclear envelope during nuclear reassembly post mitosis (Crabbe et al. 2012). Furthermore, increased mobility of dysfunctional telomeres has been observed in response to TRF2 deletion and appears to be mediated by the Linker of Nucleoskeleton and Cytoskeleton (LINC) complex (Lottersberger et al. 2015). Furthermore, induction of DSBs at telomeres using the FokI endonuclease triggered directional mobility and RAD51-dependent telomere clustering in U2OS cells (Cho et al. 2014). In both cases, no repositioning of telomeres to the nuclear periphery has been reported. Furthermore, both studies examined the impact of DSBs on telomere mobility, and thus, the effect of replication defects on nuclear relocalization of telomeres in mammalian cells remains unknown.

Here, we aimed to uncover pathways that allow the proliferation of cells expressing cancer-associated *POT1* mutations. We performed a CRISPR-interference-based genetic screen alongside a proximity ligation-based proteomic approach. Our approaches identified several synthetic lethal pathways that underscored replication stress as a vulnerability associated with POT1 dysfunction. Consistent with unresolved replication problems, we detected an increase in the frequency of mitotic DNA synthesis (MiDAS) at telomeres when POT1 was impaired. In addition, we uncovered a role for the NPC in resolving telomere replication defects in mammalian cells, and observed F-actin dependent accumulation of telomeres at the nuclear periphery in POT1 mutant cells. In summary, our study unveils a conserved function for the NPC in resolving replication defects at specialized loci from yeast to man.

## Results

### Genome-wide CRISPR interference screen identifies synthetic lethal interactions in cells expressing mutant POT1

To uncover genetic vulnerabilities in cells expressing pathogenic *POT1* mutations, we performed a synthetic lethal (SL) screen in human cells transduced with wild-type POT1 (POT1-WT), POT1-ΔOB, and POT1-K90E (Fig. S1A-B). POT1-ΔOB lacks the first OB fold and served as a surrogate for cancer-associated POT1 variants that cannot bind to ssDNA (Loayza and De Lange 2003; Hockemeyer et al. 2007; Calvete et al. 2015). The K90E substitution did not prevent DNA binding *in vitro* (Pinzaru et al. 2016), but was repeatedly identified in somatic and familial cancers (Quesada et al. 2011; Wilson et al. 2017), and triggered mild telomere dysfunction (Pinzaru et al. 2016). Both alleles retained functional interaction with TPP1 and were efficiently recruited to telomere chromatin (Loayza and De Lange 2003; Pinzaru et al. 2016).

We performed a genome-wide loss-of-function screen in human fibrosarcoma HT1080 cells using CRISPR-interference (CRISPRi) (Gilbert et al. 2014; Horlbeck et al. 2016). Cells expressing POT1-WT, POT1-ΔOB, and POT1-K90E were transduced with a catalytically inactive Cas9 fused to the KRAB domain (dCas9-KRAB) and a hCRISPRi v2.1 sgRNA library, and were passaged at ~1000x coverage for ~ 9 population doublings (PD). We performed next-generation sequencing (NGS) and determined the relative change in sgRNA abundance between PD=9 and PD=0 (Fig. 1A). Normalized sgRNA counts from two independent replicates were analyzed using the Bayesian Analysis of Gene EssentiaLity (BAGEL) algorithm to determine ‘essential’ genes (Hart et al. 2015; Hart and Moffat 2016). The likelihood of a gene depletion causing fitness defect is represented as log Bayes Factor (BF) score that is generated using empirically-determined reference gene sets (Hart et al. 2017). Using a threshold of BF ≥ 3 as a cut-off (Hart et al. 2017; Lenoir et al. 2018), we identified 1483 ‘essential’ genes in cells expressing POT1-WT. On the other hand, cells expressing POT1-K90E and POT1-ΔOB yielded a list of 1721 and 1532 ‘essential’ genes, respectively (Supplementary Table 1). Precision-recall curves confirmed the robustness of the CRISPRi screens (Fig. S1C). Furthermore, we compared the BF scores from two independent screens and observed strong correlation (r2 >0.7 - Fig. S1D and Pearson’s coefficient > 0.85 - Supplementary Table 2). To gain a better understanding of the genetic interactions with *POT1* mutations in the context of untransformed cells, we repeated the CRISPRi screen in p53-deficient retinal pigmental epithelial cells immortalized with hTERT (RPE-1). We transduced RPE-1 p53^−/−^ cells expressing dCas9-KRAB as well as POT1-WT, POT1-K90E, and POT1-ΔOB (Fig. S1B) with hCRISPRi v2.1 sgRNA library. We passaged cells for 14 PDs, then applied NGS and BAGEL analysis to determine the list of ‘essential’ genes in the different genetic conditions (Fig. 1A, Fig. S1E&F and Supplementary Table 1).

**Figure 1:**
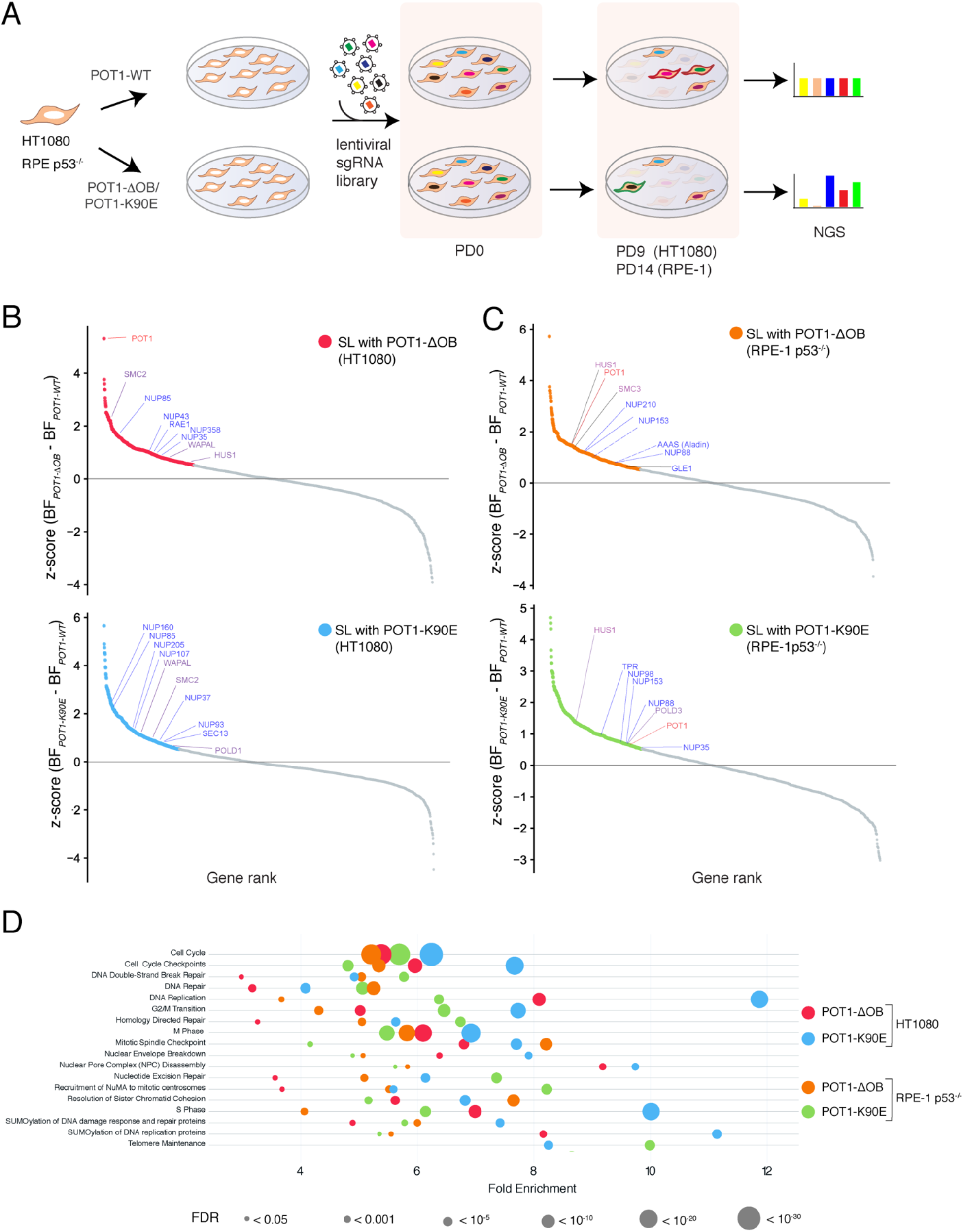
CRISPR interference screens identify synthetic lethalities in cells expressing *POT1* mutations. (**A)** Schematic of the CRIPSRi screening pipeline. HT1080 and RPE-1 p53^−/−^ cells stably expressing dCas9-KRAB, as well as POT1-WT, POT1-ΔOB and POT1-K90E, were transduced with the genome-wide hCRISPRi v2.1 sgRNA library (Horlbeck et al., 2016). Cells were collected following the selection of sgRNA-expressing cells and designated population doubling (PD) 0. Cells were also collected after ~ 9 PDs and ~ 14 PDs for HT1080 and RPE-1 p53^−/−^ cells, respectively. The relative change in sgRNA abundance at the time of final collection compared to PD0 was determined by next-generation sequencing (NGS). **(B)** Ranked z-scores of the difference in log Bayes Factor (BF) for ‘essential’ genes in POT1-ΔOB *vs.* POT1-WT (top) and POT1-K90E *vs.* POT1-WT (bottom) in HT1080 cells. BF scores were determined using the BAGEL analysis pipeline (Hart and Moffat, 2016). Genes with a z-scores ≥ 0.53 were considered to be potentially synthetic lethal (SL) with mutant POT1. SL candidates are marked in red for POT1-ΔOB and in blue for POT1-K90E. **(C)** Similar analysis as in (**B**) for RPE-1 p53^−/−^ cells. SL candidates are marked in orange for POT1-ΔOB and in green for POT1-K90E. **(B) (C)** Selected SL candidate genes highlighted in blue depict nucleoporins (NUPs), purple denotes mitotic DNA synthesis (MiDAS) and general replication stress related genes, and POT1 is highlighted in red. (**D)** Reactome pathway over-representation analysis of synthetic lethal genes with mutant POT1. The analysis was performed using PANTHER classification (v. 14.1). Fold enrichment of each pathway is plotted on the x-axis. The false discovery rate (FDR) associated with the fold enrichment is indicated by the size of the circle (the lower the FDR, the larger the circle size).

### Biological processes related to DNA replication, mitosis, and the nuclear pore complex display a genetic interaction with mutant POT1

Having confirmed the robustness of the CRISPRi screens in two independent cell lines (Fig. S1C), we then determined genes that differentially impact the growth of cells expressing mutant POT1. ‘Essential’ genes identified in cells expressing POT1-K90E and POT1-ΔOB were assigned a z-score which we calculated from the difference in BF scores between mutant and wild-type POT1. Genes with a z-score ≥ 0.53 were considered to be SL candidates (Fig. 1B&C; Supplementary Table1). The majority of candidate SL genes in RPE-1 p53^−/−^ cells expressing POT1-K90E and POT1-ΔOB were classified as ‘non-essential’ in cells expressing POT1-WT (Fig. S1F; Supplementary Table 1). Furthermore, we noted significant overlap in SL genes between POT1-K90E and POT1-ΔOB expressing cells (Fig. S2A). Notably, *POT1* appeared as an SL hit in cells expressing POT1-ΔOB, and to a lesser extent, POT1-K90E (Fig. 1B&C). This provided internal validation for our screen since a reduction in endogenous levels of POT1 with CRISPRi is expected to exacerbate telomere dysfunction caused by mutant POT1. A common SL hit that appeared in three of the four screens was *HUS1*, a member of the 9-1-1 complex that activates ATR signaling in response to replication stress (Saldivar et al. 2017). We transduced RPE-1 p53^−/−^ cells expressing dCas9-KRAB and the various POT1 alleles with sgRNAs against *HUS1* and monitored the cell growth using CellTiter-Glo and clonogenic assays. Our data corroborated the results of the CRISPRi screen, whereby HUS1 depletion conferred a greater fitness defect in cells expressing mutant POT1 (Fig. S2B&C).

We next undertook a comprehensive approach and highlighted biological pathways that were essential for growth of cells expressing *POT1* OB-fold mutations. We analyzed the SL gene lists using the PANTHER system (Mi et al. 2019) and identified processes that were common to cells expressing POT1-K90E and POT1-ΔOB. Over-representation analysis revealed an enrichment of several Reactome pathways, including G2/M transition, homology directed repair, SUMOylation of DNA repair, DNA replication proteins, and chromosome segregation during mitosis (Fig. 1D; Supplementary Table 3). These pathways could be broadly categorized as processes that mediate the cellular response to DNA replication stress, and in agreement with our previous report, firmly establish the impact of POT1 alterations on telomere replication (Pinzaru et al. 2016). Genes that fall in the “chromosome segregation during mitosis” pathway included *WAPAL* and *SMC2*, which are key factors in the mitotic DNA synthesis (MiDAS) pathway (Fig. 1B,C) (Minocherhomji et al. 2015). In addition, our screen identified the “Nuclear Pore Complex (NPC) Disassembly” pathway as essential for growth of cells with *POT1* OB-fold mutations (Fig. 1D, Supplementary Table 3). Several nucleoporins (NUPs) displayed SL interactions with POT1-K90E and POT1-ΔOB, including subunits of the Y-complex (NUP37, NUP43, NUP85, NUP107, NUP160 and SEC13) and the nuclear pore basket (NUP153 and TPR) (Fig. 1B&C).

### A proteomic approach uncovers the telomere interactome in cells expressing mutant POT1

In an independent set of experiments, we complemented our genetic screen with a proteomic approach that interrogated the changes in telomere chromatin caused by dysfunctional POT1. To do so, we applied the proximity-labelling method – termed BioID – that harnesses a promiscuous *E.coli* biotin ligase (BirA*) capable of covalently attaching biotin on proximal proteins (Roux et al. 2012). Myc-tagged BirA* was fused to POT1-WT and POT1-ΔOB and stably expressed in HT1080 cells (Fig. 2A). As a control, we showed that cells expressing BirA*-POT1-ΔOB displayed increased levels of telomere-dysfunction induced foci (TIFs), determined by the accumulation of DNA damage factor 53BP1 at telomeres (Takai et al. 2003) (Fig. S3A). Furthermore, immunofluorescence (IF) confirmed that upon treatment of cells with biotin, the fusion proteins stimulated the biotinylation of factors in the vicinity of telomeres (Fig. S3A and Fig. 2B). We then performed streptavidin immunoprecipitation (IP) on lysates from cells expressing the various POT1 alleles (Fig. S3B). Unique peptides were identified using liquid chromatography – mass spectrometry (LC-MS) and their abundance determined using intensity-based absolute quantification (iBAQ) (Schwanhausser et al. 2011) (Supplementary Table 4). We excluded proteins common to a contaminant repository for affinity purification (CRAPome) (Mellacheruvu et al. 2013) and ones recovered in <2 IPs. Based on these criteria, we identified ~200 proteins that were enriched at telomeres regardless whether they were bound by POT1-WT or POT1-ΔOB (Fig. 2C-E; Fig. S3C; Supplementary Table 4). Common hits included members of the shelterin complex – TPP1, TIN2, and TRF2 – that were recovered with similar abundance, and other previously identified telomeres associated proteins (Fig. 2D) (Dejardin and Kingston 2009; Grolimund et al. 2013; Garcia-Exposito et al. 2016). We also identified 36 proteins that were unique to, or significantly enriched (>50% in at least 2 IPs) at telomeres in the presence of POT1-ΔOB (Fig. 2E). These included 53BP1 and its interacting partners, SAP130, ARID1A and ZFR (Fig. 2E) (Gupta et al. 2018). In addition, we retrieved 13 factors that were also identified as SL candidates in the CRISPRi screen, including the MiDAS factor WAPAL and the NPC subunits, TPR and NUP153 (Fig. 1B and Fig.2E; Supplementary Tables 1&4).

**Figure 2:**
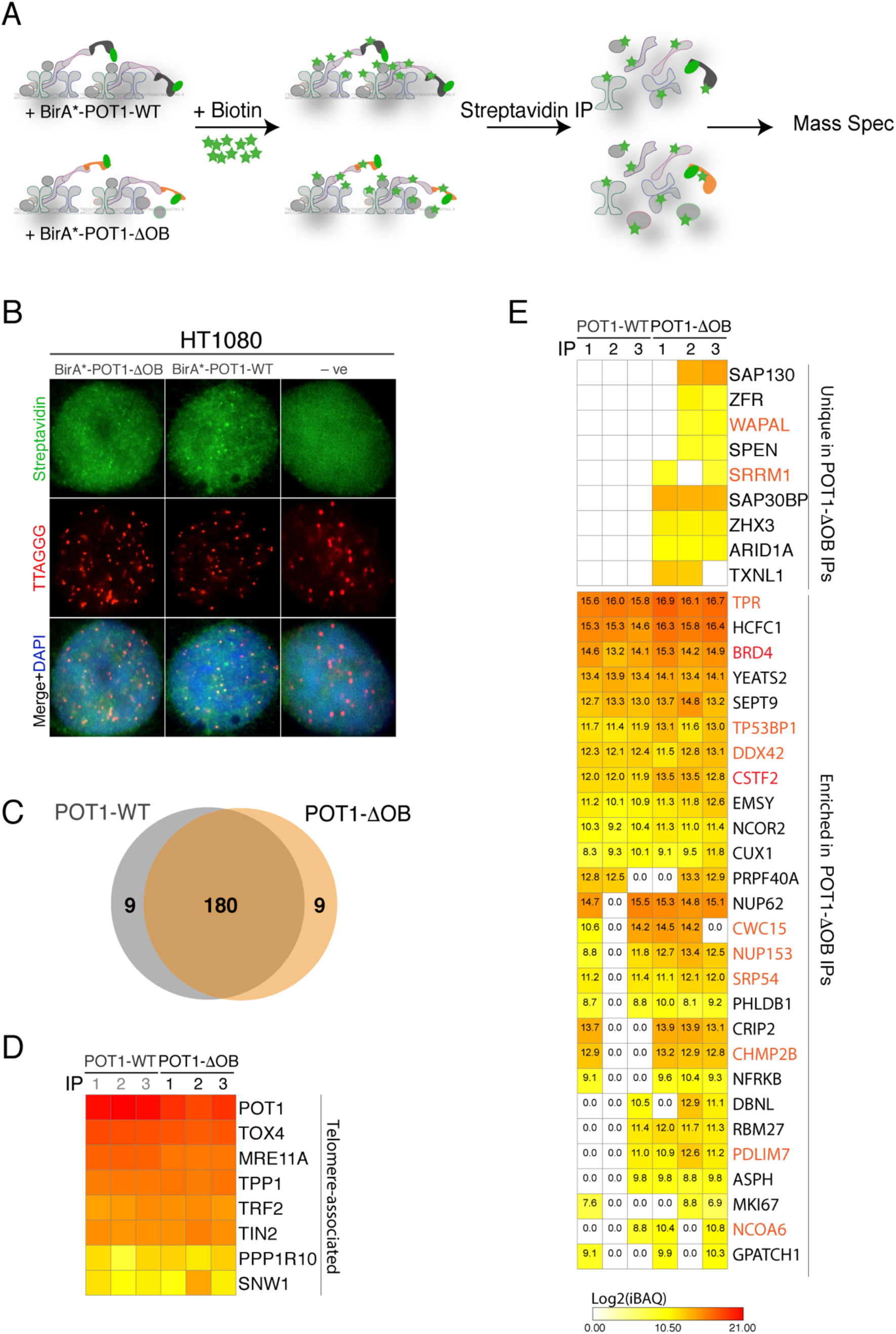
Proteomic analysis of the telomere interactome in cells expressing POT1 OB-fold mutation. **(A)** Schematic of the biotin ligase (BirA*) purification scheme to characterize the telomere proteome in cells expressing BirA*-POT1-WT and BirA*-POT1-ΔOB. BirA* covalently attaches biotin to lysine residues on vicinal proteins. Labeled proteins are recovered by streptavidin immuno-precipitation (IP) and identified using liquid chromatography - mass spectrometry. IPs were performed in triplicate for each POT1 variant. **(B)** Indirect immunofluorescence (IF) for streptavidin (green) coupled with fluorescence *in situ* hybridization for telomeres (FISH) (red) in HT0180 cells with the indicated treatment and following overnight incubation with excess biotin. Cells expressing an empty vector were used as negative control for the staining. **(C)** Venn diagram of the overlap between POT1-WT and POT1-ΔOB hits identified in ≥ 2 IPs. **(D)** Heatmap of the log2-transformed intensity-based absolute quantification (iBAQ) values for telomere-associated proteins recovered in cells expressing BirA*-POT1-WT or BirA*-POT1-ΔOB. n=3 independent IPs. **(E)** Heatmap of the log2-transformed iBAQ values of proteins enriched at telomeres in POT1-ΔOB cells. (Top) proteins uniquely present in ≥ 2 IPs in POT1-ΔOB expressing cells. (Bottom) iBAQ values for proteins with ≥ 50% enrichment (≥0.6 in log2) in ≥ 2 IPs in cells expressing POT1-ΔOB compared to those expressing POT1-WT. Values were compared in a paired manner (i.e. IP#1 fpr POT1-WT with IP#1 for POT1-ΔOB). Highlighted in red are enriched proteins that have been identified as SL candidates in the genome-wide screen (Supplementary Table1).

### Activation of Mitotic DNA synthesis (MiDAS) in response to POT1 dysfunction

When faced with incomplete DNA replication in S phase, mammalian cells use a salvage pathway termed mitotic DNA repair synthesis (MiDAS), to complete DNA copying in the early stages of mitosis (Minocherhomji et al. 2015; Bhowmick et al. 2016). Key MiDAS factors; *WAPAL*, *SMC2*, *POLD1* and *POLD3* (Minocherhomji et al. 2015; Ozer and Hickson 2018) were essential for growth of cells expressing mutant POT1 (Fig. 1B&C). Furthermore, *WAPAL* was enriched at telomeres in cells expressing POT1-ΔOB (Fig. 2E). To test whether mutations in POT1 triggered MiDAS at telomeric loci, we monitored the incorporation of base thymidine analogue EdU during mitosis (Garribba et al. 2018). In asynchronous cells, we detected a two-fold increase in the incorporation of mitotic EdU in cells expressing POT1-ΔOB compared to POT1-WT (Fig. 3A, S4A). This phenotype was also observed when cells were treated with low doses of aphidicolin followed by Cdk1 inhibition (RO-3306) to enrich for cells at the G2/M transition (Fig. 3B-D, S4B&C) (Özer et al. 2018).

**Figure 3:**
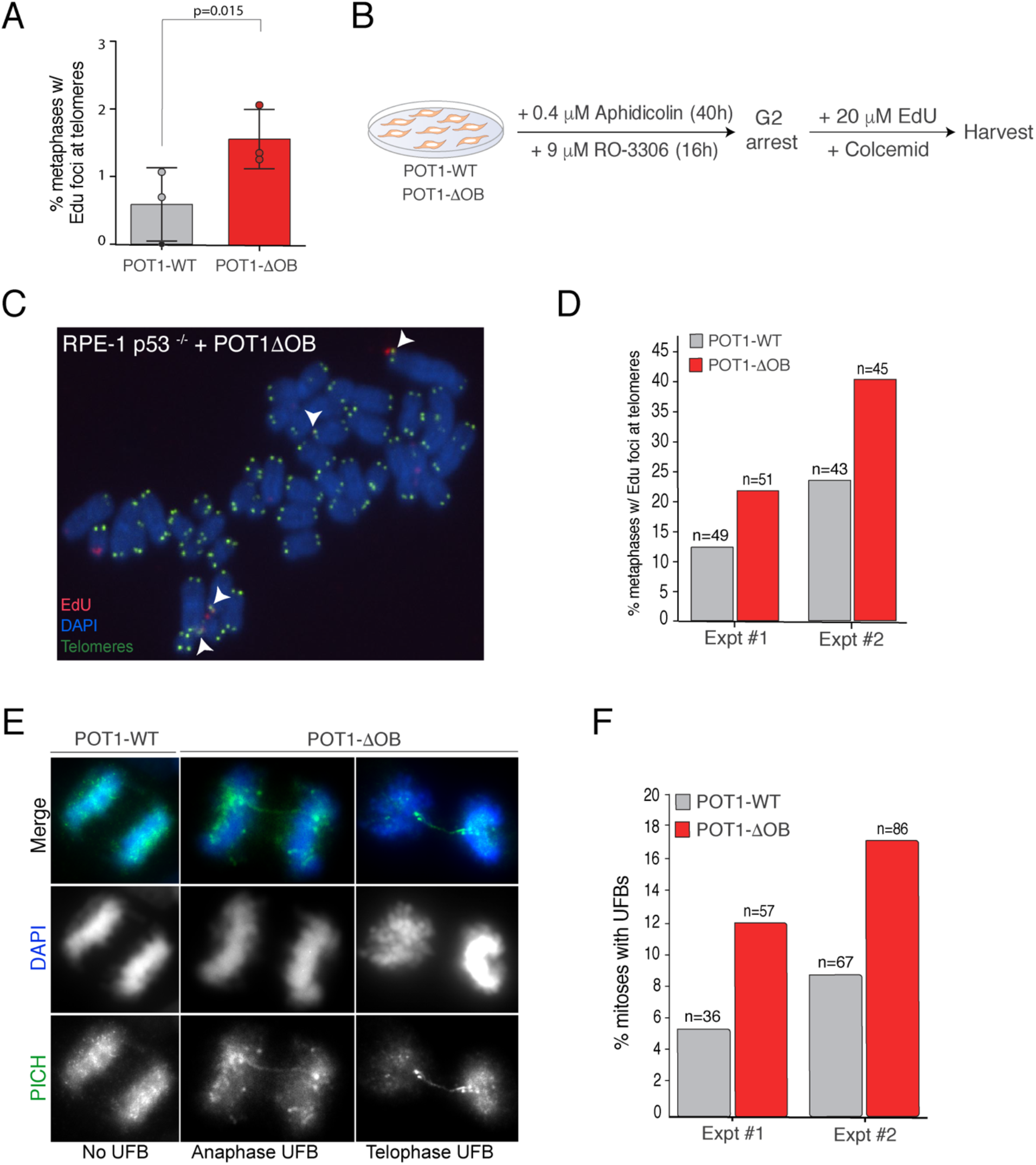
POT1 dysfunction triggers mitotic DNA synthesis (MiDAS) at telomeres. **(A)** Analysis of MiDAS in asynchronous HT1080 cells expressing POT1-WT and POT1-ΔOB. Quantification of the number of metaphases with EdU foci at telomeres from asynchronous cells. Mean ± SD of 3 independent experiments (n = 634 metaphases for POT1-WT and 597 for POT1-ΔOB) student’s t-test; paired, one-tailed. **(B)** Setup of MiDAS experiment in RPE-1 p53^−/−^ cells expressing POT1-WT and POT1-ΔOB. Cells were treated with aphidicolin (0.4 µM) for 40h and with Cdk1 inhibitor RO-3306 (9 µM) during the last 16h of the aphidicolin treatment. Following release from the G2/M block, cells were incubated with 20 µM EdU and Colcemid for 50-60 min, before metaphases were harvested. **(C, D)** Analysis of mitotic DNA synthesis (MiDAS) in RPE-1 p53^−/−^ cells exogenously expressing POT1-WT and POT1-ΔOB following the treatment as in (B). (C) Representative image of EdU incorporation at telomeres; arrowheads point at EdU foci at telomeres. (D) Quantification of the number of metaphases with EdU incorporation at telomeres; two independent experiments **(E, F)** Analysis of ultrafine DNA bridges (UFBs) in U2OS cells exogenously expressing POT1-WT and POT1-ΔOB. (E) Representative images of mitotic UFBs detected by indirect immunofluorescence for PICH (green); DNA is stained with DAPI in blue. (F) Quantification of PICH UFBs in two independent experiments (total n > 100 mitoses for each condition).

A characteristic feature of replication defect is the appearance of ultra-fine anaphase bridges (UFBs), which arise when a cell enters anaphase without having resolved replication intermediates. UFBs manifest as thin DNA entanglements typically coated with PICH (Plk1-interaction checkpoint helicase) and are observed at fragile sites (Chan et al. 2009), including telomeres (Barefield and Karlseder 2012). We examined POT1 mutant cells in mitosis and noticed an enrichment in PICH-labelled bridges compared to control cells (Fig. 3E&F). Notably, we did not detect an increase in the frequency of chromatin bridges and lagging chromosomes upon POT1 inhibition (Fig. S4D&E). Furthermore, we observed no difference in the onset of anaphase (Fig. S4F&G), indicating that replication stress in response to POT1 alterations did not lead to overt defects in cell cycle progression and chromosome segregation. Based on these results, we conclude that increased MiDAS enables POT1 mutant cells to cope with telomere replication defects, while unresolved replication intermediates that progress through mitosis give rise to UFBs.

### The NUP62 subcomplex is critical for maintaining telomere stability in cells with mutant POT1

The CRISPRi screen highlighted a genetic interaction between POT1 OB-fold mutations and several nucleoporins (NUPs) that belong to different NPC subcomplexes (Fig. 4A). In addition, TPR, NUP62, and NUP153 were enriched in the vicinity of telomeres bound by POT1-ΔOB (Fig. 2E). To uncover the function of the NPC during telomere dysfunction, we treated RPE-1 p53^−/−^ cells expressing POT1-ΔOB and POT1-WT with shRNAs against NUP62 and NUP58, independently (Fig. S5A). Assessment of cellular proliferation corroborated the results of the CRISPRi screen and showed that depletion of NPC subunits strongly impaired growth of cells expressing OB-fold mutations of *POT1* (Fig. 4B). Consistent with previous reports (Hockemeyer et al. 2007), expression of POT1-ΔOB induced a telomere DNA damage response that was exacerbated upon the depletion of NUP62 and NUP58 (Fig. 4C&D and S5B). As a control, we showed that depletion of NUP62 and NUP58 in cells expressing POT1-WT did not elicit a TIF response (Fig. 4C&D). We next examined the impact of NUP depletion on telomere fragility, which is a marker of replication stress. Chromosome analysis revealed that inhibition of NUP62 and NUP58 subunits increased the incidence of fragile telomere in cells expressing POT1-ΔOB cells (Fig. 4E&F). In conclusion, our data highlight a key role for the NPC is maintaining telomere stability in response to replication stress in human cells expressing *POT1* OB-fold mutations.

**Figure 4:**
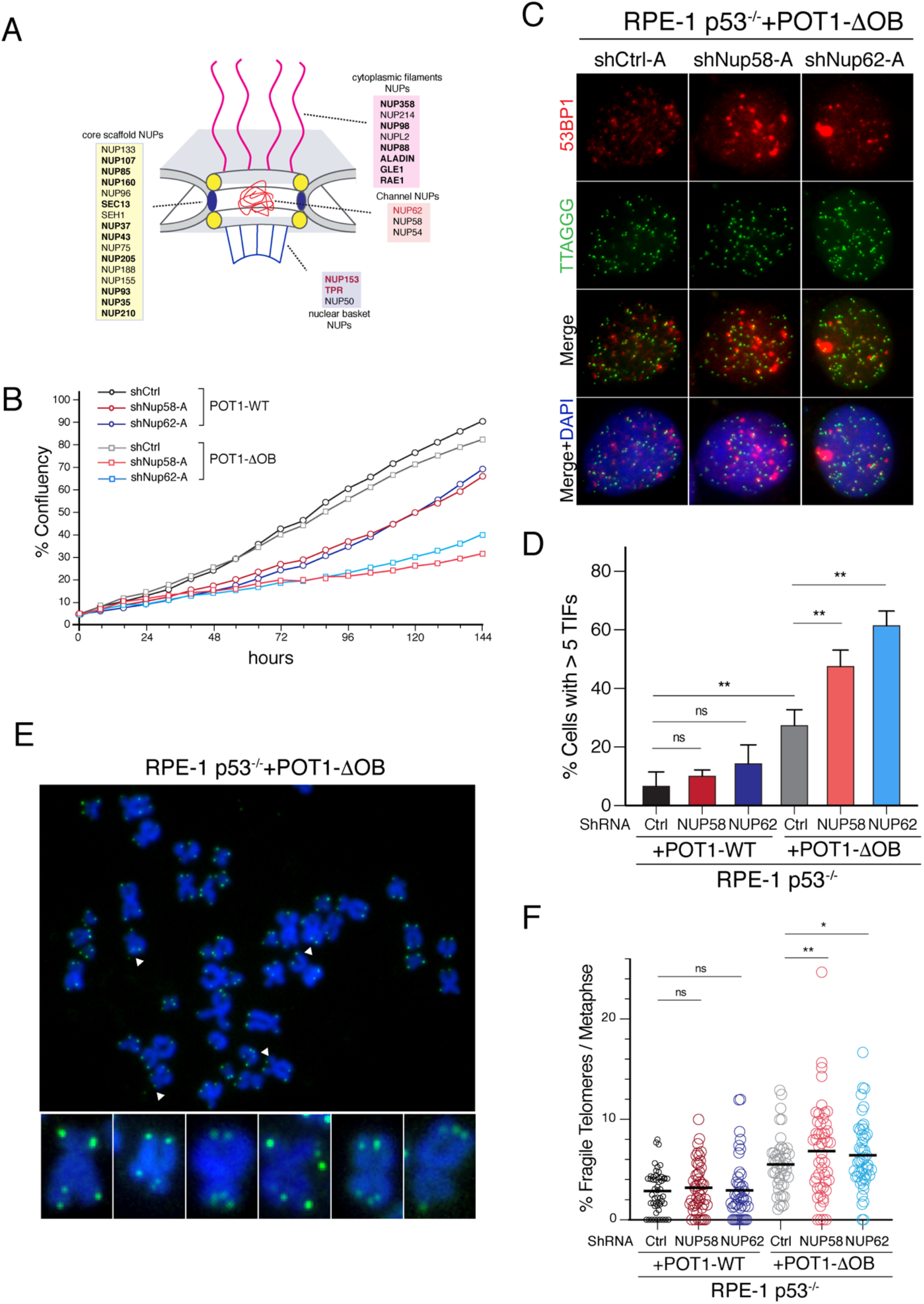
The NUP62 complex is necessary to maintain telomere integrity in cells carrying mutant POT1. **(A)** Schematic of the nuclear pore complex in mammalian cells (Kabachinski and Schwartz 2015). Highlighted in bold are SL hits with POT1 mutants and in red are nucleoporins (NUPs) enriched in the POT1-ΔOB BioID IPs. NUP153 and TPR were hits in both screens. **(B)** IncuCyte growth analysis RPE-1 p53^−/−^ cells expressing POT1-WT and POT1-ΔOB, treated with shRNAs against NUP62, NUP58, and scramble control. Cell proliferation was monitored over 160h. Graph represent data from two independent experiments. **(C)** Representative images displaying telomere dysfunction-induced foci (TIFs) in RPE-1 p53^−/−^ cells expressing POT1-ΔOB and treated with shRNAs against NUP58 and NUP62, as well as control shRNA. 53BP1 in red is detected by indirect immunofluorescence, and telomeres are marked with FISH in green. DNA is counterstained with DAPI in blue. **(D)** Quantification of the percentage of cells with 5 or more TIFs in RPE-1 p53^−/−^ cells with the indicated treatment. Graph represents the mean of n=3 independent experiments with SD (two-tailed t-test). **(E-F)** Analysis of telomere fragility in RPE-1 p53^−/−^ cells expressing POT1-WT and POT1-ΔOB and treated with the indicated shRNA. (E) Representative images showing metaphase chromosomes from POT1-ΔOB cells with fragile telomeres. Arrowheads indicate examples of fragile telomeres on the metaphase. Telomeres were stained with FISH in green, while DNA is detected with DAPI in blue. (F) Quantification of fragile telomeres per metaphase in cells with the indicated treatments. Graph represents mean of four independent experiments (n >40 metaphase per condition) with SD (one -way Annova test).

### POT1 dysfunction promotes F-actin dependent telomere localization to the nuclear periphery

So far, our results implicate the NPC in the resolution of replication-related telomere defects in human cells. This raises an obvious question related to the underlying mechanism that targets telomeres to the vicinity of nucleoporins at the nuclear periphery. A novel mechanism that mediates DNA relocalization emerged from recent literature, whereby polymerization of nuclear F-actin filaments was shown to facilitate the mobilization of damaged DNA towards the nuclear periphery. This included the directed movement of heterochromatic DSBs in *D. melanogaster* and stalled replication forks in human cells towards the nuclear envelope (Ryu et al. 2015; Caridi et al. 2018; Lamm et al. 2018; Schrank et al. 2018). To investigate telomere targeting to the nuclear periphery, we visualized the three-dimensional telomere localization in cells harboring mutant POT1. Specifically, RPE-1 p53^−/−^ cells expressing POT1-ΔOB and POT1-WT were transfected with a GFP-actin chromobody that also express a nuclear localization signal (NLS). Cells were then fixed, stained with anti-TRF2 antibody, and visualized in three-dimensions using AiryScan super-resolution microscopy. Marking the nuclear area with diffuse GFP signal enabled us to segment the nucleus into six zones of equal volumes (Fig. 5B) and assign telomeres to different volumetric zones (Fig. 5A&B and Fig. S6A). We observed that telomeres were randomly distributed throughout the nucleoplasm in cells expressing POT1-WT. In contrast, telomeres in POT1-ΔOB expressing cells were enriched towards the nuclear edge (Fig. 5A&B and Fig. S6A). We then treated cells expressing POT1-ΔOB and POT1-WT with Latrunculin B (LatB) to suppress F-actin polymerization and observed a reduction in peripheral localization of telomeres in cells with mutant POT1 (Fig. 5B and Fig. S6B). In conclusion, our data highlight telomere repositioning to the nuclear periphery in response to replication stress and hint at a possible role for nuclear F-actin in this process.

**Figure 5:**
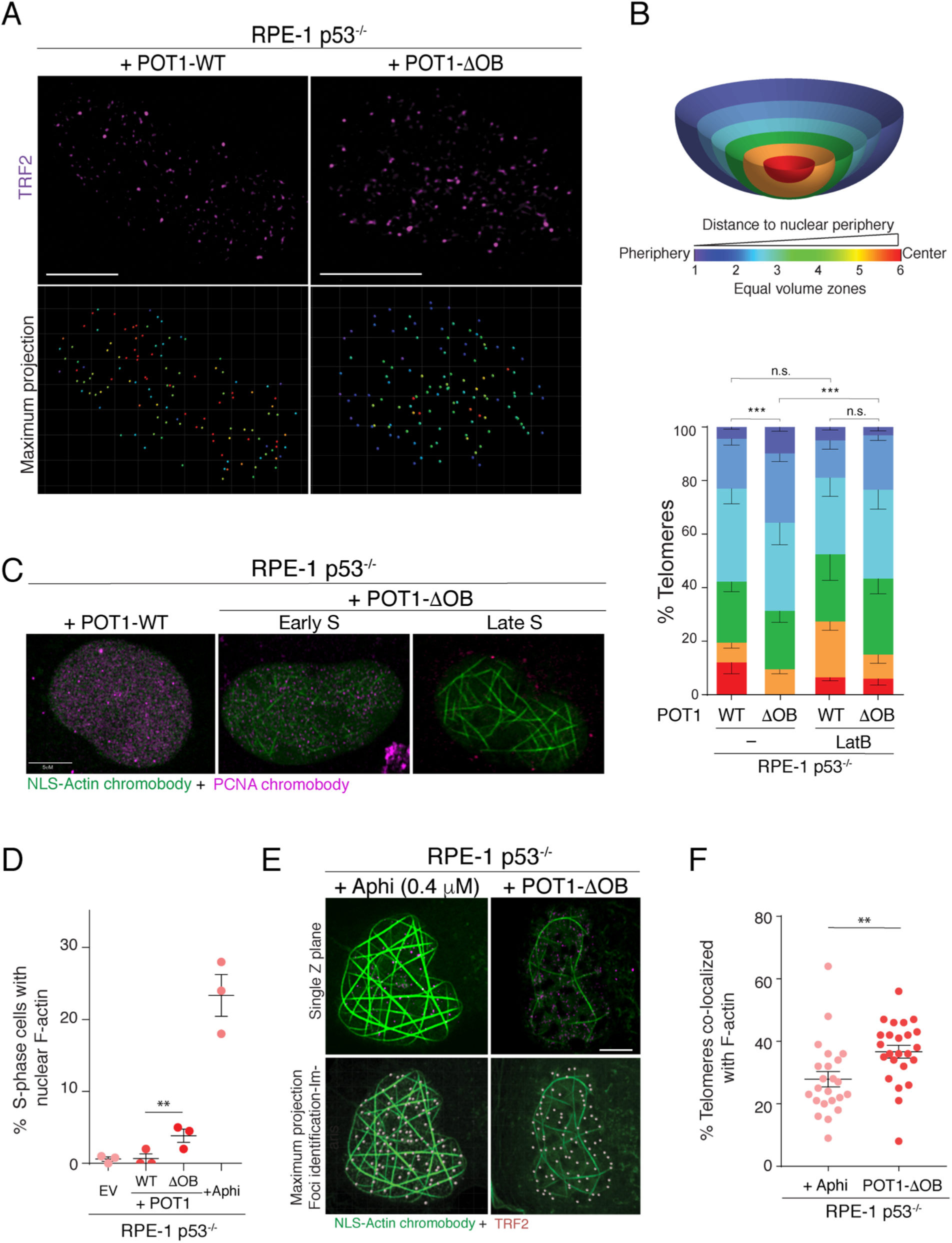
Nuclear F-actin polymerization in POT1-ΔOB cells facilitate the relocalization of telomeres to the nuclear periphery. **(A)** Representative super-resolution microscopy of a three-dimensional (3D) image through the nuclear volume of fixed RPE-1 p53^−/−^ cells expressing POT1-WT and POT1-ΔOB. Telomeres were detected with an anti-TRF2 antibody. **(B)** Quantification of telomere localization in the nucleus of cells imaged in (A), in the absence or presence of 0.2 µM Latrunculin B (LatB) treatment (24h prior to fixation).Top panel: Telomeres were identified throughout the nuclear volume and their distance to the nuclear periphery calculated using the Imaris 8.4.1 software. Nuclei were segmented into six equal volume zones from nuclear center to the periphery and each telomere assigned to the corresponding zone. The zone for each telomere relative to the nuclear periphery is identified via color coding. Lower panel: Graph represents distribution of telomeres with respect to the nuclear periphery. Mean ± SEM (chi-square test). n ≥ 1805 telomeres from >19 nuclei and 3 independent experiments. **(C)** Representative super-resolution microscopy of single Z-planes taken from 3D images through the nuclear volume of fixed RPE-1 p53^−/−^ cells expressing POT1-WT and POT1-ΔOB. Cells were transfected with NLS-GFP-Actin and TagRFP-PCNA chromobodies 48h prior to fixation. (**D)** Quantification of filamentous-actin (F-actin) positive S-phase nuclei from the images depicted in C). Each data point represents an individual biological replicate. n=3 independent experiments with > 168 nuclei analyzed per experiment. SEM with Fisher’s exact test. **(E)** Representative super-resolution microscopy of 3D images through the nuclear volume of fixed parental RPE-1 p53^−/−^ cells and cells expressing POT1-ΔOB. Cells were transfected with an NLS-GFP-Actin chromobody 48h prior to fixation and telomeres were detected with an anti-TRF2 antibody. Parental cells were treated with 0.4 µM aphidicolin for 24h prior to fixation. **(F)** Quantification of the percentage of telomeres that co-localized with nuclear F-actin in experiments highlighted in (E). For all panels: **p < 0.001, ***p < 0.0005. Scale bar represents 5 µM. n=3 independent experiments with > 23 nuclei and >1656 telomeres. SEM with Mann-Whitney test.

Lastly, we assayed cell cycle-dependent nuclear F-actin polymerization upon telomere dysfunction. Cell expressing POT1 variants were labelled with an NLS-GFP-actin chromobody that detects nuclear F-actin filaments and co-stained with an RFP-PCNA chromobody to mark cells in S-phase (Fig. 5C). As a control, we showed that treatment of wild-type cells with low levels of aphidicolin (0.4 µM), which causes global replication stress, triggered significant nuclear F-actin filament formation (Fig. 5D). Furthermore, S-phase cells displaying nuclear F-actin were enriched in cells expressing POT1-ΔOB compared to POT1-WT(Fig. 5C&D). Albeit, the induction of nuclear F-actin in the presence of POT1-ΔOB was less robust than aphidicolin treatment. Telomere DNA represents a small fraction of the entire genome which could explain why telomere-specific replication stress results in a milder effect on nuclear F-actin polymerization (Fig. 5D). Importantly, the colocalization of telomeres with nuclear F-actin filaments was significantly enriched in cells expressing POT1-ΔOB relative to those treated with aphidicolin (Fig. 5E-F). It is worth noting that the colocalization of telomeres with F-actin in cells expressing POT1-WT could not be analyzed due to the paucity of S-phase cells with nuclear actin filaments. Taken together, our data are consistent with a model where actin-dependent polymerization in cells undergoing replication stress due to *POT1* OB-fold mutations promotes telomere repositioning to the nuclear periphery where NUPs reside.

## Discussion

### Replication stress as a major vulnerability in cells with mutant POT1 alleles

Sequencing of cancer genomes identified POT1 mutations in several solid tumors and hematological malignancies (Quesada et al. 2011; Lazzerini-Denchi and Sfeir 2016; Speedy et al. 2016; McMaster et al. 2018). To better understand the basis by which mutant POT1 alleles foster tumorigenesis, we performed a genome-wide CRISPRi screen. Our screen identified a set of pathways that were essential for survival of cells with *POT1* OB-fold mutations. These pathways included DNA replication, S phase, homologous recombination, SUMOylation of DNA replication proteins, and nuclear-pore complex disassembly (Fig. 1D), and collectively underscore replication stress as a major defect associated with *POT1* mutations. While nuclease-based CRISPR screens often yield stronger SL hits, a CRISPR interference-based approach has an advantage as it uncovers genetic interactions that involve essential genes. Indeed, the identification of SL interactions between mutant POT1 and subunits of the NPC, which are essential for cellular survival (Kabachinski and Schwartz 2015), was possible because of the incomplete repression of genes by CRISPRi (Gilbert et al. 2014).

We complemented our genetic screen with a biotin-dependent labeling approach (Roux et al. 2012) and identified factors that are in proximity to telomeres bound by mutant POT1. Notable telomere interactors that were enriched in POT1-ΔOB expressing cells and also appeared as SL candidates in the CRISPRi screen included WAPAL and subunits of the NPC (Fig. 2). Consistent with POT1 dysfunction leading to incomplete telomere replication in S phase, we detected an increase in MiDAS in cells expressing POT1-ΔOB (Fig. 3). These results highlight components of MiDAS as potential targets for the treatment of an increasing number of tumors associated with *POT1* mutations.

### A conserved function for the NPC in resolving damage in repetitive DNA

The link between the nuclear pore complex and DNA repair was first established in yeast based on the observation that mutations in several NUPS cause hypersensitivity to DNA damage agents and are lethal with mutations that impair HR (Bennett et al. 2001; Loeillet et al. 2005; Therizols et al. 2006). Since then, increasing evidence demonstrated the relocalization of persistent DNA breaks to the vicinity of nuclear pores in *S. cerevisiae*. Such lesions included collapsed replication forks, HO-induced breaks, eroded telomeres, and CAG repeats (Nagai et al. 2008; Khadaroo et al. 2009; Oza et al. 2009; Su et al. 2015; Churikov et al. 2016). In all cases, relocalization to the nuclear pore was necessary for efficient DNA repair and to prevent genome instability. Transient tethering of damaged DNA to the nuclear pore was also noted in *Drosophila melanogaster* cells, particularly at breaks incurred within heterochromatic loci (Ryu et al. 2015). With regards to mammalian cells, no direct role for the NPC in repairing DNA lesions has been reported thus far. Here, we provide the first genetic evidence that links the NPC with maintenance of repetitive DNA and demonstrate targeting of damaged telomeres to the nuclear periphery.

Mouse telomeres rendered dysfunctional as a result of TRF2 deletion displayed increased mobility but did not cluster at the nuclear periphery (Lottersberger et al. 2015). Similarly, FokI-induced DSBs at telomeres in U2OS cells triggered telomere clustering without apparent relocalization to the periphery (Cho et al. 2014). In contrast, our data suggest that telomeres undergoing replication stress have distinct properties that promote their targeting to nuclear periphery (Fig. 5A&B). Telomere uncapping upon TRF2 loss and FokI induced telomere breaks trigger a canonical DSB response that activates the ATM kinase pathway (Celli and de Lange 2005) (Doksani and de Lange 2016). In contrast, stalled forks as a result of *POT1* inhibition activate the ATR kinase pathway (Pinzaru et al. 2016), potentially implicating this S phase specific kinase in telomere repositioning to the nuclear periphery. Though, it is also possible that telomere targeting to the periphery is driven by a specific type of substrate that is associated with stalled replication, such as a reversed or collapsed fork.

### Nuclear F-Actin filaments facilitate telomere repositioning to the nuclear periphery

Repositioning of DNA breaks away from their primary site is common to several genomic loci and has been observed in many organisms (Lemaitre et al. 2014; Ryu et al. 2015; Caridi et al. 2018; Smith et al. 2018; Marnef et al. 2019). The mechanistic understanding of damage-induced DNA mobility has just started to emerge. Genetic studies in yeast and flies identified a critical role for SUMOylation, mainly by the Slx5-Slx8 SUMO-targeted ubiquitin ligase (STUbL), in the spatial and temporal regulation of break repair (Nagai et al. 2008; Ryu et al. 2015; Su et al. 2015; Churikov et al. 2016; Horigome et al. 2016). Interestingly, “SUMOylation of DNA damage response and repair proteins” and “SUMOylation of DNA replication proteins” appeared as pathways that are synthetic lethal with mutant POT1 (Fig. 1D) and the Slx5/8 human ortholog *RNF4* showed a genetic interaction with POT1-ΔOB (Supplementary Table 1). While RNF4 has been previously linked to DSB repair at telomeres in the context of TRF2 loss (Groocock et al. 2014), its function in response to telomere replication defects remains to be investigated.

Recent reports implicated nuclear F-actin filament formation during the directional mobility of damaged DNA in *D. melanogaster* as well as human cells (Caridi et al. 2018; Lamm et al. 2018; Schrank et al. 2018). Our data are consistent with nuclear F-actin polymerization underlying the process of telomere relocalization to the periphery. Specifically, we observed F-actin filament formation in S phase cells expressing mutant POT1 (Fig. 5 C-D) and show that blocking actin polymerization with LatB reduced the fraction of telomeres at the nuclear periphery (Fig. 5B). Based on these data, we propose a model where actin polymerization potentially facilitates the targeting of replication-defective telomeres to the NPC. The relocalization event is expected to isolate stalled forks involving telomeric repeats, presumably to prevent illegitimate recombination repair and facilitate fork restart (Freudenreich and Su 2016). In summary, our study uncovered a conserved mechanism from yeast to man that targets persistent damage at specialized loci to the nuclear periphery to ensure genome stability.

## Acknowledgements

We acknowledge Alireza Khodadadi-Jamayran for assistance with bioinformatic analysis and Michael J. Smith for comments on the manuscript. We thank the Proteomics core at NYU School of Medicine (The mass spectrometric experiments were in part supported by NYU School of Medicine and the Laura and Isaac Perlmutter Cancer Center Support grant P30CA016087 from the National Cancer Institute). We acknowledge the microscopy core at NYUSOM for help with image analysis. We thank the UCSF Center for Advanced Technology for help with the NGS. This work was supported by a grant from the NIH-NCI (U01CA231019) to A.S.

## Authors Contribution

A.S., E.L-D and A.M.P. conceived the experimental design. A.M.P performed experiments with help from M.A. N.L. performed experiments related to Figure 5., under the guidance of A.J.C. A.S. and A.M.P. wrote the manuscript. All authors discussed the results and commented on the manuscript.

## Authors information

Agnel Sfeir is a co-founder, consultant, and shareholder in Repare Therapeutics. Correspondence and requests for materials should be addressed to A.S.

## MATERIALS AND METHODS

### Cell culture

RPE1-hTERT *TP53*^−/−^ were a gift from the Meng-Fu Bryan Tsou Lab at Memorial Sloan Kettering Cancer Center. RPE1-hTERT *TP53*^−/−^ and osteosarcoma U2OS cells were cultured in DMEM supplemented with 10% FBS (Gibco), 2 mM L-glutamine (Sigma), 100 U μg/ml penicillin (Sigma), 0.1 μg/ml streptomycin (Sigma) and 0.1 mM non-essential amino acids (Invitrogen). HT1080 fibrosarcoma cells were grown in DMEM supplemented with 10% BCS (bovine calf serum, Hyclone), 2 mM L-glutamine (Sigma), 100 U μg/ml penicillin (Sigma), 0.1 μg/ml streptomycin (Sigma) and 0.1 mM non-essential amino acids (Invitrogen). The cells were tested for mycoplasma using the LookOut® Mycoplasma PCR Detection Kit (Sigma, # MP0035), following the manufacturer’s instructions and using the JumpStart™ Taq DNA Polymerase (Sigma, #D9307).

### Plasmids

Human POT1 variants (wild-type POT1, POT1-ΔOB, POT1-K90E) were subcloned into the lentiviral construct pHAGE2-EF1a-MCS-IRES-blast and pWZL-N-Myc-IRES-hygro from pLPC-N-Myc-POT1. The pHAGE2-EF1a-MCS-IRES backbone was a gift from the Matthias Stadtfeld lab at NYU Langone Health. Wild-type POT1 and POT1-ΔOB were also subcloned downstream of Myc-FLAG-BirA* to generate pLPC-N-Myc-FLAG-BirA*-POT1/POT1-ΔOB constructs.

The hCRIPSRi-v2 library top 5 sgRNAs/ gene (Horlbeck et al. 2016)(Addgene # 83969), was a gift from the Jonathan Weissman lab at UCSF. The lentiviral construct expressing dCas9-HA-NLS-TagBFP-Krab-NLS was a gift from the Jonathan Weissman lab. We generated a dCas9-HA-NLS-mCherry-Krab-NLS construct by replacing the TagBFP with mCherry, using *BamHI* and *NdeI* sites added on mCherry during PCR amplification with the Q5® High-Fidelity DNA Polymerase (NEB, #M0491). The pBabe-H2B-GFP plasmid (Addgene #26790) was a gift from the Susan Smith lab at NYU Langone Health. The constructs were transduced into human cells using established transduction protocols for generating stable cell lines.

### CRISPRi screen: infection, cell culture, library preparation

The screen was performed with the hCRIPSRi-v2 library (Horlbeck et al. 2016), which comprises 5 optimized sgRNAs per gene, for a total of 102640 sgRNAs targeting close to 19000 genes. The library also includes ~1900 non-targeting control sgRNAs. The hCRIPSRi-v2 is a two-vector system, where the dCas9-KRAB fusion is expressed separately from the sgRNA library. The screen was performed in duplicate in HT1080 cells expressing dCas9-HA-NLS-BFP-Krab-NLS and pWZL-Myc-POT1-IRES-Hygro, pWZL-Myc-POT1-K90E-IRES-Hygro or pWZL-Myc-ΔOB-POT1-IRES-Hygro. The whole-genome screen was repeated once in RPE1-hTERT *TP53*^−/−^ cells transduced with dCas9-HA-NLS-mCherry-Krab-NLS and POT1 variants expressed in the pHAGE2-EF1a-MCS-IRES-blast vector. The experiment was performed following guidelines described before (Gilbert et al. 2014). Briefly, the sgRNA library was transfected into HEK293T cells together with 3^rd^ generation lentiviral packaging vectors to generate lentivirus, the viral supernatant was collected at 48h and 72 h, snap frozen and stored at −80°C until use. After determining the transduction efficiency in each cell line, a large-scale transduction was performed aiming to achieve a low multiplicity of infection (MOI) (achieved MOI of 0.3-0.4 for HT1080, MOI of ~0.6 for RPE1-hTERT *TP53*^−/−^) and a coverage of >300x of the library. After two days from the infection, the cells were selected with puromycin to enrich for the sgRNA integration, then the final representation was determined. HT1080 cells were selected with puromycin 800 ng/mL for 2 days, while the RPE1-hTERT p53 null cells were selected with 20 µg/mL puromycin for 4 days. Following selection, at least 2×125 mil. cells were frozen down per condition as time point 0 samples (T0), while the rest of the cells were further cultured, plating the equivalent number of cells to achieve ~1000x coverage of the library per condition. The cells were split every 2-3 days, aiming to maintain 1000x coverage of the library at plating throughout the screen. HT1080 cells were cultured in maintenance hygromycin (50 µg/mL), while RPE1-hTERT p53 null cells were cultured in maintenance blasticidin (7.5 −10 µg/mL) to ensure the exogeneous expression of the POT1 constructs is maintained throughout the screen.

The endpoint was reached after ~ 9 population doublings for HT1080 cells and ~14 population doublings for the RPE1-hTERT p53 null cells. At least 2×125 mil. cells were frozen down per condition at the endpoint. Following genomic extraction, the DNA was digested overnight with ~400 U/mg SbfI-HF (New England Biolabs, Ipswich, MA) in order to enrich for the cassette containing the sgRNA construct. Subsequently, sgRNAs were amplified by PCR and, during the enrichment, Illumina adapters were added to the 3’ and 5’ end of the sgRNA cassette together with an index barcode at one end. Following PCR cleanup, the samples were sequenced on an Illumina HiSeq-4000 using custom primers, single-end 50 base pairs with a 6bps index. For each analyzed condition, we obtained at least ~ 50 million total reads per condition.

Custom sequencing primers:

**5’ Sequencing Primer:**

5’ GTGTGTTTTGAGACTATAAGTATCCCTTGGAGAACCACCTTGTTG

**3’ Sequencing Primer**;

5’ TGATAACGGACTAGCCTTATTTAAACTTGCTATGCTGTTTCCAGCTTA

### Bioinformatic analysis of the CRISPRi screen

Sequencing reads were aligned to the library sequences using a custom Python script (available at https://github.com/mhorlbeck/ScreenProcessing). For the HT1080 analysis the sgRNA read counts were normalized to account for differences in total reads across samples. Then, the normalized reads were averaged across the two replicates per condition at each time point. The averaged read counts were used to compute the fold changes at the endpoint compared to day 0 for all conditions. sgRNAs with < 25 reads at the T0 timepoint were excluded from the fold-change calculation. Subsequent analysis of the screen was done using the Bayesian Analysis of Gene EssentiaLity (BAGEL) algorithm (Hart et al. 2015; Hart and Moffat 2016). Empirically-determined reference gene sets, including 687 core essential genes and 927 non-essential genes (Hart et al. 2017) were used to generate log2 Bayes Factor scores. The BAGEL algorithm was ran using bootstrapping and the network boosting option, which uses the preexisting information from the functional associations annotated in the STRING v10.5 database (Szklarczyk et al. 2017) to refine the log2 Bayes Factor scores. Log2 Bayes Factors (BF) were also computed for the individual HT1080 replicates, starting from normalized read counts, and using the cross-validation option for BAGEL. The correlation between the HT1080 replicates for each condition was determined with Pearson’s coefficient and r-squared analysis. The performance of the screen was determined using the reference sets of essential and nonessential genes to compute precision-recall curves. To determine the different fitness genes between the wild-type and mutant POT1 conditions, a z-score reflecting a ‘differential essentiality’ score was computed (Steinhart et al. 2017) for each ‘essential’ gene (BF>3) in the mutant POT1 condition: for each analyzed gene, the BF score for POT1-WT was subtracted from the BF score for the respective POT1 mutant; z-scores were subsequently assigned to each BF scores difference, in order to highlight the most ‘differential essential’ genes between the wild-type and mutant POT1 conditions. The analysis for RPE1-hTERT p53 null cells was done similarly as for HT1080, using the BAGEL algorithm employing cross-validation and the network boosting option.

Pathway enrichment analysis was conducted using PANTHER 14.1 (http://www.pantherdb.org/) (Mi et al. 2019) with the default statistical overrepresentation test parameters for Reactome pathways (Fabregat et al. 2018).

Data was plotted using matplotlib vs 3.1.1.

### BirA*-mediated biotinylation and pull down of proteins

HT1080 cells stably expressing empty vector, pLPC-N-Myc-FLAG-BirA*-POT1 or pLPC-N-Myc-FLAG-BirA*-POT1-ΔOB were grown in 15 cm^2^ dishes to 90% confluence at the time of harvesting. Approximately 100 million cells per condition were treated with biotin (50 µM) for 16 – 20 hours prior to harvesting. The cells were then collected by trypsinization, pooled and counted. The cells were the pelleted, washed twice with 1x PBS and resuspended in cold NP40 lysis buffer (10mM Tris-HCl pH 7.4; 10mM NaCl; 3mM MgCl2; 0.5% NP40 with freshly added protease inhibitors (Roche)). In order to allow for a gentle fractionation, the resuspended cells were incubated rocking for ~ 1h @ 4°C, until an enriched nuclear fraction could be observed under the light microscope. The nuclei were pelleted 10 min @ 4°C, 3300g, resuspended very well in room temperature SDS lysis buffer (1% SDS; 10mM EDTA pH 8; 50mM Tris-HCl pH8, with freshly added 1mM PMSF and protease inhibitors), then incubated 10 min on ice. Next, the samples were boiled 5 min @ 95°C. After cooling on ice, the samples were sonicated on Biorupter UCD-200 ice water bath 10 sec ON/ 20 sec OFF, HIGH setting until they became clear and fluid (at least 10 min). Subsequently, the samples were centrifuged 10min @ 13000 rpm, 4°C to clear any debris. The supernatant was diluted 1:10 with IP Dilution Buffer (1.2mM EDTA; 16.7mM Tris-HCl pH8; 150mM NaCl with freshly added 1 mM PMSF and protease inhibitors), loaded on Amicon Ultra 3K Columns and spun 30 min @ 4000g, 4°C to remove free biotin and concentrate the sample. The concentrate was transferred into fresh tubes and supplemented with Triton X-100 to 1% final concentration. The amount of concentrate used per IP was equalized among samples based on the initial cell number calculated during harvest. After saving ~ 5% of lysate as input, the rest was incubated overnight with Dynabeads® MyOne™ Streptavidin C1 (ThermoFisher, #65001) @ 4°C, rotating end-over-end. The next day, the beads were washed 2 x Buffer 1 (2% SDS in water) @ RT, 1 x Buffer 2 (0.1% deoxycholate; 1% Triton X-100; 500mM NaCl; 1mM EDTA; 50mM HEPES pH7.5)@ 4°C, 1 x Buffer 3 (250mM LiCl; 0.5% NP40; 0.5% Deoxycholate; 1mM EDTA; 10mM Tris pH 8.1) @ 4°C and 2x Buffer 4 (50mM Tris-HCl pH 7.4; 50mM NaCl)@ 4°C. All washed were performed for 8-10 min using an end-over-end rotator. After the last wash, the beads were resuspended in Buffer 4 and 4X Laemmli buffer and boiled for 5 min @ 95°C to elute the bound proteins. Approximately 5% of the eluate was used for silver stain and western blot analysis, while the rest was submitted for mass spectrometry. The silver stain was done according to the manufacturer’s instructions (Pierce Silver Stain Kit # 24612, Thermo Fisher). The experiment was conducted independently three times.

### Preparation of Samples for Mass Spectrometry

The samples were resuspended in NuPAGE® LDS Sample Buffer (Novex). The proteins were reduced with 2μl of 0.2M dithiothreitol (Sigma) for one hour at 57 °C at pH 7.5. Next, the proteins were alkylated with 2μl of 0.5M iodoacetamide (Sigma) for 45 minutes at room temperature in the dark. The samples were loaded on a NuPAGE® 4-12% Bis-Tris Gel 1.0 mm (Life Technologies) and run for 8 minutes at 200V. The gel was stained with GelCode Blue Stain Reagent (Thermo). The gel bands were excised, cut into 1mm^3^ pieces and destained for 15 minutes in a 1:1 (v/v) solution of methanol and 100mM ammonium bicarbonate. The buffer was exchanged and the samples were destained for another 15 minutes. This was repeated for another 3 cycles. The gel plugs were dehydrated by washing with acetonitrile, and further dried by placing them in a SpeedVac for 20 minutes. 250ng of sequencing grade modified trypsin (Promega) was added directly to the dried gel pieces followed by enough 100mM ammonium bicarbonate to cover the gel pieces. The gel plugs were allowed to shake at room temperature and digestion proceeded overnight. The digestion was halted by adding a slurry of R2 50 μm Poros beads (Applied Biosystems) in 5% formic acid and 0.2% trifluoroacetic acid (TFA) to each sample at a volume equal to that of the ammonium bicarbonate added for digestion. The samples were allowed to shake at 4°C for 120 mins. The beads were loaded onto C18 ziptips (Millipore), equilibrated with 0.1% TFA, using a microcentrifuge for 30 s at 6,000 rpm. The beads were washed with 0.5% acetic acid. Peptides were eluted with 40% acetonitrile in 0.5% acetic acid followed by 80% acetonitrile in 0.5% acetic acid. The organic solvent was removed using a SpeedVac concentrator and the sample reconstituted in 0.5% acetic acid.

### Mass Spectrometry Analysis

An aliquot of each sample was loaded onto an Acclaim PepMap trap column (2 cm x 75 µm) in line with an EASY-Spray analytical column (50 cm x 75 µm ID PepMap C18, 2 μm bead size) using the auto sampler of an EASY-nLC 1000 HPLC (Thermo Fisher Scientific) with solvent A consisting of 2% acetonitrile in 0.5% acetic acid and solvent B consisting of 80% acetonitrile in 0.5% acetic acid. The peptides were gradient eluted into an Orbitrap Elite Mass Spectrometer (Thermo Fisher Scientific) using the following gradient: 5 - 35% in 60 min, 35 - 45% in 15 min, followed by 45 - 100% in 5 min. The gradient was held at 100% for another 10 minutes. MS1 spectra were recorded with a resolution of 60,000, an AGC target of 1e6, with a maximum ion time of 200ms, and a scan range from 400 to 1500m/z. Following each full MS scan, fifteen data-dependent MS/MS spectra were acquired. The MS/MS spectra were collected in the Ion Trap with an AGC target of 3e4, maximum ion time of 150ms, one microscan, 2m/z isolation window, dynamic exclusion of 30 sec, and Normalized Collision Energy (NCE) of 35.

### Data Processing for Mass Spectrometry

The MS/MS spectra were searched against the UniProt *Homo sapiens* reference proteome database (downloaded 10/2015) using Andromeda (Cox et al. 2011). Protein quantitation was performed using Intensity-based absolute quantification (iBAQ)(Schwanhäusser et al., 2011) within MaxQuant (Cox and Mann 2008) (Version 1.5.3.30). The mass tolerance was set to 20ppm for MS1. MS2 searches were set to a tolerance of 0.5Da. False discovery rate (FDR) filtration was done first on peptide level and then on protein level. Both filtrations were done at 1% FDR using a standard target-decoy database approach. All proteins identified with less than two unique peptides were excluded from analysis.

### Guide RNA sequences

sgA _HUS1: 5’ GAGCCGCGGCGGGCCTCTGT

sgB_HUS1: 5’ GTACCCACAGAGGCCCGCCG

sgC_HUS1: 5’ GGGATGAAGGCGGCTTTCAA

non-targeting-sgA: 5’ GCGTACGACAATACGCGCGA

non-targeting-sgB: 5’ GTTACTACAGGAGCCGAAGG

non-targeting-sgC: 5’ GGCGCCGGACTGGACCTCGA

non-targeting-sgD: 5’ GGCGCTCCCACCGATAAAGT

### shRNA sequences

The shRNAs against *NUP58 (NUPL1)* and *NUP62* were obtained from the MISSION shRNA Sigma-Aldrich library:

shRNA_NUP58-A: CCGGGCTTCAGTTTAGGATTCAATACTCGAGTATTGAATCCTAAACTGAAGCTTTTT

shRNA_NUP58-B: CCGGGCGGCACAACTTCAGTCTATTCTCGAGAATAGACTGAAGTTGTGCCGCTTTTT

shRNA_NUP62-A: CCGGGCAACCTCACTAATGCCATATCTCGAGATATGGCATTAGTGAGGTTGCTTTTT

shRNA_NUP62-B: CCGGGCTTTCATTTGAGTATCTTTGCTCGAGCAAAGATACTCAAATGAAAGCTTTTT

Scramble control: CCGGCCTAAGGTTAAGTCGCCCTCGCTCGAGCGAGGGCGACTTAACCTTAGGTTTTT

### Cell Titer Glo assay

SgRNAs targeting *HUS1* (A, B, C) and non-targeting guides RNAs (A, B, C, D) from the CRISPRi library individually cloned into lentiGuide-puro were used to virally transduce RPE-1 p53 null cells expressing dCas9-HA-NLS-mCherry-Krab-NLS and POT1 variants cloned into pHAGE2-EF1a-MCS-IRES-blast. At 48h post-infection, the cells were put on puromycin selection (20ug/mL) to enrich for the sgRNA-expressing cells. Following completion of the puromycin selection, the cells were trypsinized, counted twice and following serial dilutions, 500 cells were plated in triplicate per condition in 96 wells. The cells were left to grow until they neared confluence in the non-targeting guides wells (8 days for POT1-WT/ POT1-ΔOB; 7 days for POT1-K90E), then cell growth was quantified using CellTiter-Glo (Promega) according to the manufacturer’s instructions. The growth was normalized to the average growth in the non-targeting sgRNA-A wells for each cell line (POT1-WT, POT1-K90E, POT1-ΔOB respectively).

### Clonogenic survival assay

SgRNAs targeting *HUS1* (A, B) and non-targeting guides RNAs (A, B) from the CRISPRi library individually cloned into lentiGuide-puro were used to virally transduce RPE1-hTERT p53 null cells expressing with dCas9-HA-NLS-mCherry-Krab-NLS and POT1 variants cloned into pHAGE2-EF1a-MCS-IRES-blast. At 48h post-infection, the cells were trypsinized, counted twice and following serial dilutions, 500 cells were plated in duplicate on 100 mm plates per condition. The high transduction efficiency (at least 80%) was confirmed in parallel by selecting half of the remaining cells with puromycin (20 μg/mL) for 5 days and comparing the number of surviving cells in the selection condition to the number of unselected cells. After 14 days, the colonies were stained with a crystal violet staining solution (crystal violet 0.05% w/v; 1% Formaldehyde, 1x PBS, 1% methanol) for 20 min at RT, washed extensively and left to dry overnight. The colonies were counted using an in-house macro for ImageJ.

### IncuCyte® S3 Live-Cell Analysis

RPE1-hTERT p53 null cells expressing either POT1-WT or POT1-ΔOB cloned into pHAGE2-EF1a-MCS-IRES-blast were transduced with pLKO.1-puro scramble control, shRNA_NUP58 or shRNA_NUP62. Following puromycin selection, 25000 cells per condition were plated in 12 wells and 16 fields were imaged per well every 8h using the 10x objective on the IncuCyte® S3 live-cell analysis system (Essen Bioscience Inc.). After 160h, when the control cells were nearing confluence, we used the IncuCyte® Analysis Software with the default settings and the 1.2 segmentation adjustment to determine the % of confluence over time in each well. For the cleanup step, the hole fill was set to 0 and no adjustments were made to the pixel size. No area or eccentricity filters were applied. Two independent experiments were performed.

### Mitotic DNA Synthesis (MiDAS) detection and telomeric FISH

Detection of mitotic DNA synthesis on metaphase plates was performed according to the experimental procedures for detecting MiDAS on metaphase chromosomes published by (Garribba et al. 2018). To note, HT1080 cells expressing exogenous POT1 variants were not synchronized or treated with aphidicolin for MiDAS. HT1080 cells were incubated with 20 µM EdU and 100 ng/mL colcemid (Roche) for 75 min to arrest cells in metaphase. RPE1-hTERT p53 null cells expressing exogenous POT1 variants were treated with 0.4 µM aphidicolin for 40h. In the last 16h of the aphidicolin treatment the RPE1-hTERT p53 null cells were concurrently treated with 9 µM RO3306 to enrich for cells in G2/M. The RPE1-hTERT p53 null cells were washed 3x with 1xPBS to release them from arrest, then incubated for 50-60 min at 37^ο^C with 20 µM EdU and 100 ng/mL colcemid (Roche). Following metaphase arrest, the cells were harvested, incubated in freshly made hypotonic 0.075 M KCl solution at 37^ο^C for 20-30 min, followed by an overnight fixation in cold methanol/acetic acid (3:1). Metaphase spreads were generated by dropping cells onto pre-chilled glass slides. EdU detection was performed with a Click-IT EdU Alexa Flour 594 Imaging kit (Life Technologies), as described previously (Garribba et al. 2018). Telomeric FISH was performed as previously described (Sfeir et al. 2009) using an Alexa488-labeled TelC PNA probe (PNA Bio Inc.). The cell cycle profiles of untreated vs. aphidicolin and RO3306 treated RPE1-hTERT p53 null cells were determined using propidium iodide staining.

### Detection of Ultrafine Anaphase Bridges

Ultrafine anaphase bridges in U2OS osteosarcoma cells exogenously expressing wild-type POT1 and POT1-ΔOB were detected using the mouse monoclonal anti-PICH antibody (clone 14226-3; Millipore, 04-1540; 1:100 dilution), by following the co-extraction protocol described by (Bizard et al. 2018)

### Live cell imaging

RPE1-hTERT p53 null cells stably expressing exogenous POT1 variants and H2B-GFP were plated on uncoated 35 mm Glass Bottom Microwell dishes (MatTek, No. 1.5 coverglass, P35G-1.5-14-C) and incubated overnight. Approximately 6h before imaging, the media was replaced with a 1:10 ratio of regular cell culture media to Live Cell Imaging Media (Live Cell Imaging Solution (Life Technologies # A14291DJ) supplemented with 10% FBS, 2 mM L-glutamine, 100 U μg/ml penicillin, 0.1 μg/ml streptomycin, 0.1 mM non-essential amino acids, 1mM sodium pyruvate (Life Technologies #11360070) and 20 mM dextrose (Sigma # D9434). One experiment was imaged on a Nikon Eclipse Ti total internal reflection fluorescence/epifluorescence inverted microscope fitted with an Andor Zyla camera, using a Plan Fluor 40x DIC M N2 objective, 100 ms exposure and 2×2 binning. Two subsequent, independent experiments were imaged on a Nikon Eclipse Ti inverted microscope equipped with an Andor Neo camera, using a Plan Fluor 40x DIC M N2 objective, 220 ms exposure and 2×2 binning, 7% intensity of the 488 nm laser. Both microscopes were equipped with environmental chambers and automated stages. All experiments were imaged for 15 h, at 2 min intervals, capturing nine z-stacks with 1.2 µm steps. Movies were analyzed using the Nikon NIS-Elements software. Mitotic duration was determined between nuclear envelope breakdown and anaphase. Counts were blinded.

### IF-FISH

IF-FISH experiments were performed as described before (Takai et al. 2003). The primary antibodies used were 53BP1 (100-304A, rabbit polyclonal; Novus Biologicals; 1:1000), Myc (9B11, Cell Signaling; 1:1000). The secondary antibodies used were raised against rabbit or mouse and conjugated with Alexa 488 (Invitrogen) or Rhodamine Red-X (Invitrogen), and diluted 1:1000. Telomere FISH was performed using Alexa 488 or Cy3-OO-TelC labeled PNA probes (PNA Bio Inc.). Streptavidin−Alexa 488 (Invitrogen #S32354, 1:2000 dilution) was used for detection of biotinylated proteins by IF.

### Western blots

Cells were lysed in lysis buffer (50 mM Tris-HCl – pH 7.4, 150 mM NaCl, 1 mM EDTA, 0.1% SDS, 1% Triton-X, 1mM DTT, 1 mM PMSF, and supplemented with complete protease inhibitor mix (Roche),) for 30 min on ice, followed by centrifugation at maximum speed for 10 min at 4°C. An equal volume of 4x Laemmli buffer was added to the supernatants and the samples were boiled for 5 min. Approximately 1×10^5^ cells were separated on an SDS-PAGE gel, followed by blotting to nitrocellulose membranes. The membranes were blocked in 5% BSA in TBST, incubated with primary antibodies, followed by an incubation with horseradish peroxidase (HRP) secondary antibodies (Mouse IgG HRP Linked Whole Ab-GE Healthcare, #NA931, 1:5000; Rabbit IgG HRP Linked Whole Ab-GE Healthcare, #NA934, 1:5000) in 3% milk in TBST. Primary antibodies: M2 Flag (Sigma Aldrich, 1:10,000 dilution), anti-c-Myc antibody clone 9E10 (Millipore, 1:1000 dilution), anti-HA.11 clone 16B12 (Biolegend, previously Covance Catalog# MMS-101P, 1:1000 dilution) and gamma-tubulin clone GTU-88 (Sigma Aldrich, 1:5000 dilution). Streptavidin-HRP (Invitrogen, #1953050) was used for detection of biotinylated proteins.

### Imagining for telomeric localization

48 hours after transfection of the chromobody plasmids and 24 hours after addition of aphidicolin or Latrunculin B, cells were fixed in 3.7% formaldehyde/1x PBS for 10 min, then permeabilized with 0.5% Triton/1x PBS and blocked with 5% Bovine Serum Albumin (Sigma-Aldrich)/1x PBS. Slides were incubated for 3 hours at 37º C with polyclonal anti-TRF2 antibody, washed four x 5 min in 1x PBS, then incubated for one hour at 37º C with goat polyclonal anti-rabbit Alexa Fluor 647. First and secondary antibodies were diluted in 1x PBS supplemented with 5% FCS. Before mounting, slides were washed again as described above and dehydrated in a graded series (70, 90, 100%) of ethanol solutions for two min each. Slides were then mounted with Prolong Gold (Life Technologies) and imaged using super resolution microscopy.

Super resolution imaging was performed on a ZEISS LSM 880 AxioObserver confocal fluorescent microscope fitted with an Airyscan detector using a Plan-Apochromat 63x 1.4 NA M27 oil objective. Cells were imaged using 1.9% excitation power of 568 nm laser, 2% excitation power of 488 nm laser, and 1.8% excitation power of 647 nm laser, with 1×1 binning for all laser conditions in combination with the appropriate filter sets. Ten or more z stacks (167 nm) were captured with frame scanning mode and unidirectional scanning. Z-stacks were Airyscan processed using batch mode in Zen software.

Imaging data was imported into Imaris 8.4.1 software, where nuclei were segmented as “cells” using the Cell function on the actin-NLS channel. Telomeres were segmented as “vesicles” using the Cell function. The distance between foci and the nuclear periphery were calculated using the function ‘Distance of vesicles to cell membrane’.

